# Ultra-high field imaging reveals increased whole brain connectivity underpins cognitive strategies that attenuate pain

**DOI:** 10.1101/802306

**Authors:** Enrico Schulz, Anne Stankewitz, Anderson M Winkler, Stephanie Irving, Viktor Witkovsky, Irene Tracey

## Abstract

We investigate how the attenuation of pain with cognitive interventions affects the strength of cortical connections by pursuing a whole brain approach. While receiving tonic cold pain, 20 healthy participants were asked to utilise three different pain attenuation strategies. During a 7T fMRI recording, participants were asked to rate their pain after each single trial. We related the trial-by-trial variability of the attenuation performance to the trial-by-trial functional connectivity of the cortical data.

Across all conditions, we found that a higher performance of pain attenuation was predominantly associated with higher functional connectivity. Of note, we observed an association between low pain and high connectivity for regions that belong to the core areas of pain processing, i.e. the insular and cingulate cortices. For one of the cognitive strategies (safe place), the performance of pain attenuation was explained by diffusion tensor imaging metrics of increased white matter integrity.

**Impact Statement:** In a single trial analysis, the more effective attempts to attenuate pain in three different conditions are associated with general higher functional connectivity across the entire brain.

## INTRODUCTION

An increased perception of pain is generally associated with increased cortical activity; this has been demonstrated in a number of brain regions and processes involved in sensory, emotional, cognitive, and affective aspects of pain (Tracey, Irene and Mantyh, 2007; Wiech et al., 2008). Given the threatening nature of pain, the information processed from these different aspects have to be integrated and assessed to compute an appropriate decision and subsequent action (Wiech and Tracey, 2013). To do so, pain-processing brain regions are required to exchange information, which has been found to entail increased functional connectivity between relevant cortical and subcortical regions (Sprenger et al., 2015; Villemure and Bushnell, 2009). Conversely, less is known about connectivity changes during decreased pain, although many studies highlight decreased neuronal activity with some studies highlighting selective changes in coupling between brain regions (Ploner et al., 2011).

Only a few studies have investigated the network activity of the pain system by quantifying the covariation of the fluctuating blood-oxygen-level dependent (BOLD) activity between brain regions. Changes of this covariation of cortical signals have then been linked to conditions that represent different levels of pain experience. Villemure & Bushnell (2009) and Ploner et al. (2011), for example, investigated the influence of different levels of emotion and attention on pain-related cortical connectivity (Ploner et al., 2011; Villemure and Bushnell, 2009). Both studies observed an increase of connectivity for the conditions that increased the intensity of pain; i.e. increased attention towards painful stimuli was associated with more negative emotions.

A further study found that a change in pre-stimulus cortical connectivity patterns from the anterior insula to the periaqueductal grey (PAG), which is part of the descending pain modulatory system (Sprenger et al., 2018), determined whether a subsequent nociceptive stimulus was perceived as painful or not (Ploner et al., 2010). Supporting that observation, other investigations have similarly reported increased functional connectivity between the PAG and the perigenual anterior cingulate cortex (pACC) for conditions associated with decreased pain intensity perception (placebo, shift of attention) (Eippert et al., 2009; Sprenger et al., 2012; Tracey, Irene et al., 2002). Functional connections might be supported by the structural integrity of white matter tracts between brain regions, as measured using diffusion tensor imaging (DTI). A recent study showed that the strength of the descending pain modulatory system was significantly correlated to the effectiveness in alleviating pain through transcranial direct current stimulation brain stimulation (Lin et al., 2017).

Therefore, all studies to date point to the relevance of connectivity patterns in pain modulation; yet, excluding an increased connectivity to the descending pain modulatory system’s PAG, the precise nature of cortical connectivity during decreased pain is unclear and limited. Using ultra-high field functional magnetic resonance imaging (fMRI) to provide enhanced signal-to-noise ratio (SNR) to facilitate single-trial analysis, we explored the functional connections that contribute to the attenuation of pain by means of three different cognitive interventions: (a) a non-imaginal distraction by counting backwards in steps of seven; (b) an imaginal distraction by imagining a safe place; and (c) reinterpretation of the pain valence (cognitive reappraisal). These cognitive strategies are hypothesised to be represented in the brain by a complex cerebral network that connects a number of brain regions, where:

1. The effective use of a cognitive strategy that is successful for pain attenuation results in an increase of functional connectivity between task-related brain regions.
2. Decreased connectivity is expected between cortical areas that are involved in the processing and encoding of pain intensity, e.g. sub-regions of the insular cortex, the cingulate cortex, somatosensory cortices, and PAG.
3. Increased connectivity is hypothesised for the descending pain control system, particularly for the connection between the pACC and the PAG.
4. Divisions of the insular cortex and their connections to frontal and somatosensory regions play a key role through their high relevance in integrating sensory information.

Unlike previous research paradigms, the present experimental procedure aims to approximate clinical treatment procedures by using a pain stimulation approach that produces longer lasting pain experiences. Healthy participants were asked to utilise cognitive strategies in order to attenuate the experience of pain during 40s of cold stimulation. We pursued a whole-brain parcellation approach (Glasser et al., 2016) in order to assess every cortical connection that contributes to pain relief.

## RESULTS

Behavioural data revealed a significant reduction of perceived pain for all three interventions compared to the unmodulated pain condition. The most effective condition in attenuating pain intensity and pain unpleasantness was the “counting backwards” task (21±2.55 SE for intensity attenuation and 19±2.87 SE for unpleasantness attenuation; p<0.05). More detailed results are reported in our previous publication (Schulz et al., 2019).

Overall, we found an increase of connectivity during pain attenuation: as a consequence of utilising the strategies, trials utilising the cognitive tasks have been rated as less painful and had a stronger connectivity compared with trials of the unmodulated pain condition. Therefore, trials at higher levels of pain are coupled with low connectivity, and trials at lower levels of pain are coupled with high connectivity.

We pursued a whole-brain approach by subdividing the cortex into 180 regions per hemisphere plus 11 subcortical regions (Glasser et al., 2016) and related cortical connectivity to pain ratings at single trial level. This approach was facilitated by an increased SNR as a result of ultra-high field recording, as well as by a more reliable assessment of single trial data from longer lasting painful stimulation and an extended task application. For each of the three conditions, we merged the trials of the cognitive interventions with the unmodulated pain trials, which has two major advantages:

i. First, it takes the within-subjects variable performance of the pain attenuation attempts into account; e.g. a more effective attempt to attenuate pain is considered to cause a different cortical connectivity than a less effective attempt.
ii. Second, we also take into account the more natural fluctuation of the unmodulated pain trials.

The findings are represented in confusion matrices, depicting the pain intensity-related connectivity between all brain regions. Some of the regions were previously interpreted as task-related and others as belonging to pain circuits (see Schulz et al., 2019). In the present study:

Positive relationships (*red*) show connectivities that were increased in particularly effective trials (performance encoding). For all tasks, we confirmed our first hypothesis by showing that an increased connectivity of *task-processing* brain regions (see (Schulz et al., 2019) is related to particularly effective attempts to attenuate pain. In addition, we found increased brain activity for more effective single trials also in *pain-processing* regions. Negative relationships (*blue*) represent cortical connections that are disrupted in effective trials to attenuate pain. Disruptions were hypothesised to occur for the *core regions of pain processing*, such as for the various subregions of the insular, cingulate, and somatosensory cortices. However, we found that these regions predominantly showed increased connectivity during effective trials of pain attenuation (see above).

### (1) Counting

We aimed to detect patterns of connectivity changes that are related to effective trials during counting. The attenuation of pain during counting is predominantly related to an increase of cortical connectivity in several brain regions, with the exception of decreased connections involving the right temporo-parieto-occipital junction (Figure 1A). The detailed matrix of statistical results can be found in the supplementary material (Supplementary Spreadsheet 1).

**Figure 1.**
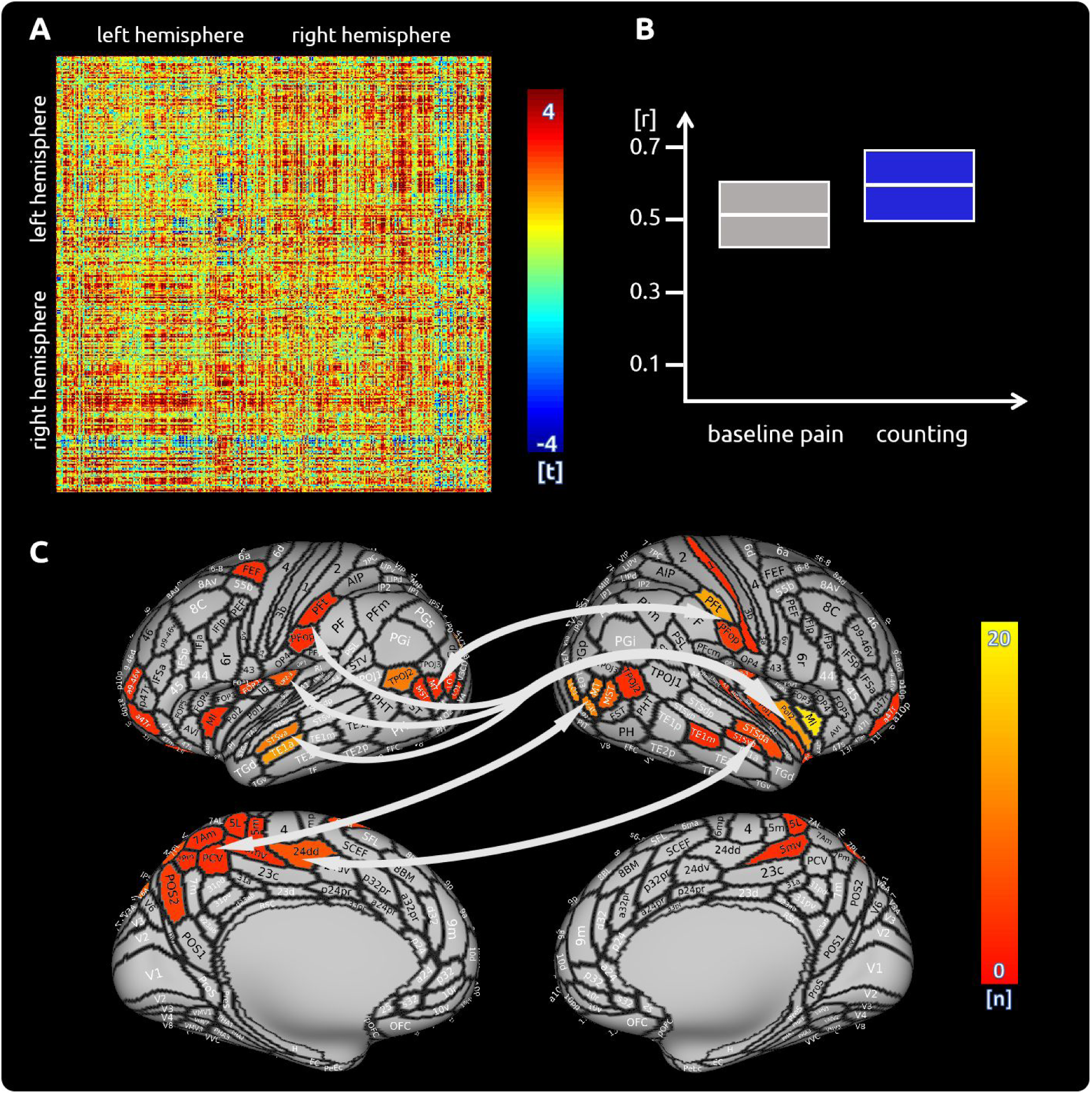
Counting: (A) the confusion matrix shows the statistical results. Each line represents one of the 371 ROIs. The values are mirrored along the principal diagonal of the matrix. A single red dot represents the varying connectivity between two specific brain regions and indicates that a stronger cortical connectivity in a single trial is related to a decrease in pain perception (performance encoding). These findings are the result of the higher connectivity in the trials of the counting task compared to unmodulated pain trials. (B) data from the confusion matrix averaged across all subjects, connections and trials (mean ± standard deviation; for illustration purposes only). (C) Depiction of the cortical regions as defined by the Glasser parcellation; The arrows show the best connected regions; the right middle insular cortex has the most connections where connectivity changes are shown to significantly modulate pain intensity. Only regions with at least 3 significant connections (n>2) are included in the cortical map. For more detailed information on the exact connections see the supplementary spreadsheets.

We found that some regions show a particularly strong connectivity: the right anterior and posterior insula, the left and right temporal cortices, the left parietal cortex, as well as higher order visual regions in occipito-temporal areas. The best connected area is the right middle insula, showing the involvement of a number of subregions (Figure 1C). Indeed, some prominent connectivity patterns are noticeable: pain attenuation-related connections from the right insular sub-regions are always connected to insular sub-regions from the contralateral hemisphere, but not to other ipsilateral insular regions. In addition, areas in the left medial wall of the parietal cortex (Brodmann area 7) are functionally connected to a right posterior cortical region that stretches from higher order visual areas (lateral occipital cortex) to the posterior medial temporal cortex. The homologue left occipito-temporal region is functionally connected to the right inferior parietal lobe (subregions PFt and PFop). Regions in the left superior and middle temporal cortex are strongly connected with several sections of the insular cortex. Extended regions in the left superior parietal cortex (Brodmann area 5) and the posterior cingulate cortex are functionally connected with the right middle insular cortex (Figure 1C, Supplementary Figure 1, Supplementary Spreadsheet 1).

Measures of structural connectivity (DTI fibre tracking) did not explain interindividual differences in modulating task-related functional connections in the counting condition.

### (2) Safe place

During the imagining condition, we found an increase of connectivity across all cortical regions when compared to the unmodulated pain condition (Figure 2A). The detailed matrix of statistical results can be found in the supplementary material (Supplementary Spreadsheet 2). There is no negative relationship between single trial connectivity and pain intensity. Besides the well-connected right insular cortex, with several significant sub-regions, we observed attenuation-related connectivity changes in right parietal (BA 5) and left superior parietal cortices (BA 7). Further well-connected areas include a frontal language area (BA 55b) as well as motor and premotor areas. Regions in the right posterior insular cortex are connected to the left parietal cortex (BA 7). The right precentral areas are functionally interconnected with prefrontal and orbitofrontal areas, the right parietal cortex (BA 5), and the left superior parietal cortex (BA 7). The right “belt” regions are functionally connected to prefrontal and orbitofrontal areas (Figure 2C, Supplementary Figure 2).

**Figure 2.**
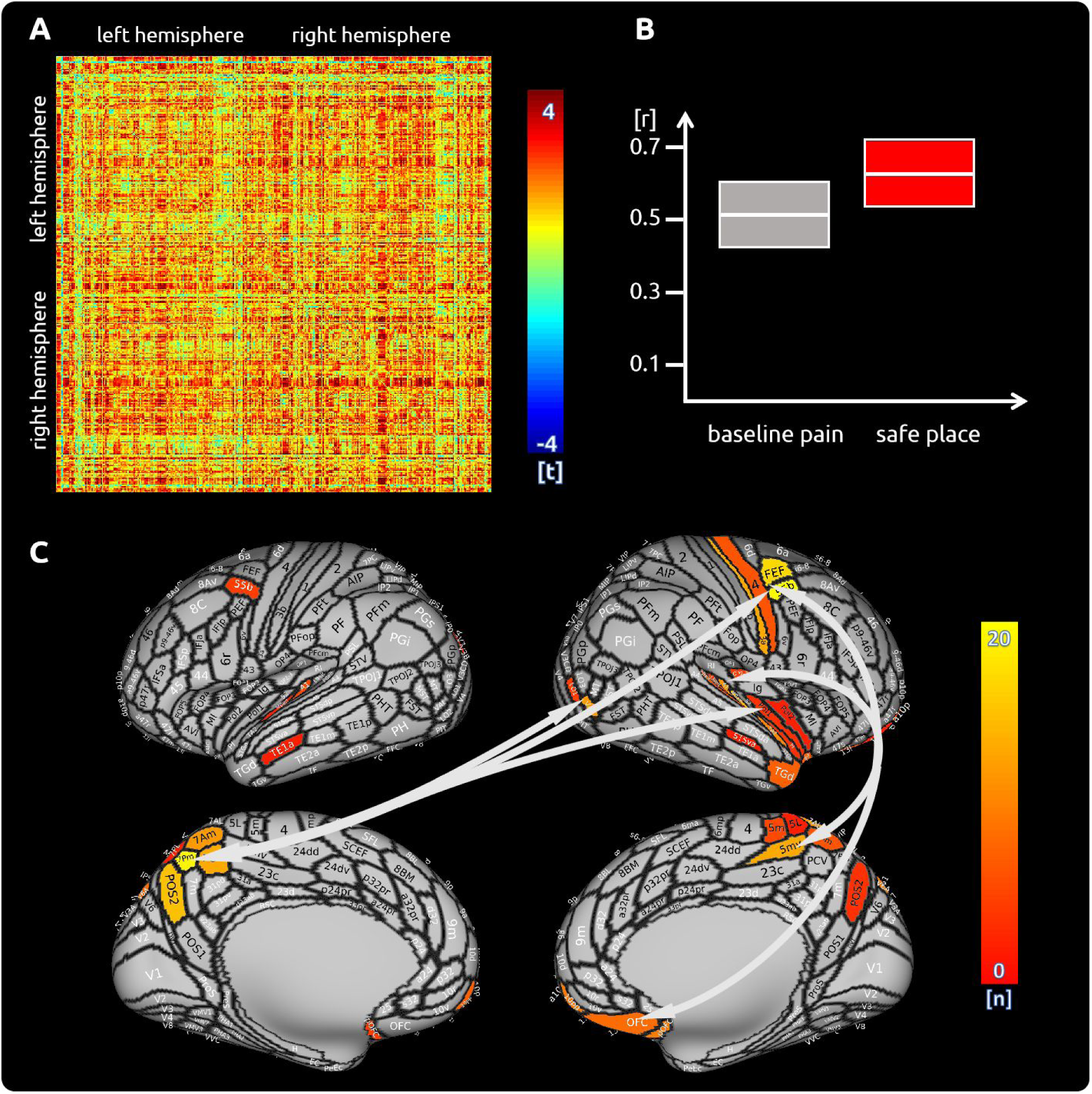
Safe place: (A) the confusion matrix shows the statistical results. Each line represents one of the 371 ROIs. The values are mirrored along the principal diagonal of the matrix. A single red dot represents the varying connectivity between two specific brain regions and indicates that a stronger cortical connectivity in a single trial is related to a decrease in pain perception (performance encoding). These findings are the result of the higher connectivity in the trials of the imagination task compared to unmodulated pain trials. (B) data from the confusion matrix averaged across all subjects, connections and trials (mean ± standard deviation; for illustration purposes only). (C) Depiction of the cortical regions as defined by the Glasser parcellation; The arrows show the best connected regions; the left parietal cortex and right premotor areas have the most connections where connectivity changes are shown to significantly modulate pain intensity. Only regions with at least 3 significant connections (n>2) are included in the cortical map. For more detailed information on the exact connections see the supplementary spreadsheets.

For the safe place condition only, we found that the strength of fibre connections mediates the strength of the functional connectivity. Some subjects made use of their better structural connectivity, as measured by the number of streamlines obtained from fibre tracking. Strong structural connectivities are related to a better ability to modulate the functional connectivity in order to attenuate pain. This applies especially to connections between frontal regions (IFSp and Brodmann area 8C) and the secondary somatosensory cortex (SII). Further functional connections that are supported by the strength of fibre connections projected to memory-related areas (presubiculum of the hippocampus and entorhinal cortex).

### (3) Reappraisal

While executing cognitive reappraisal we found a pain attenuation-related increase of functional connectivity compared to the unmodulated pain condition across the entire cortex (Figure 3A). The detailed matrix of statistical results can be found in the supplementary material (Supplementary Spreadsheet 3). Decreased functional connectivity has not been observed. Connections that included frontal premotor and insular sub-regions contributed to a decrease of pain (Figure 3C, Supplementary Figure 3). However, the main hub of connectivity was located in the medial parieto-occipital cortex. Besides other regions, the area V6A is interconnected with several insular and frontal premotor areas, some of which control eye movements. The structural characteristics between cortical regions did not contribute to an enhanced functional connectivity for reappraisal.

**Figure 3.**
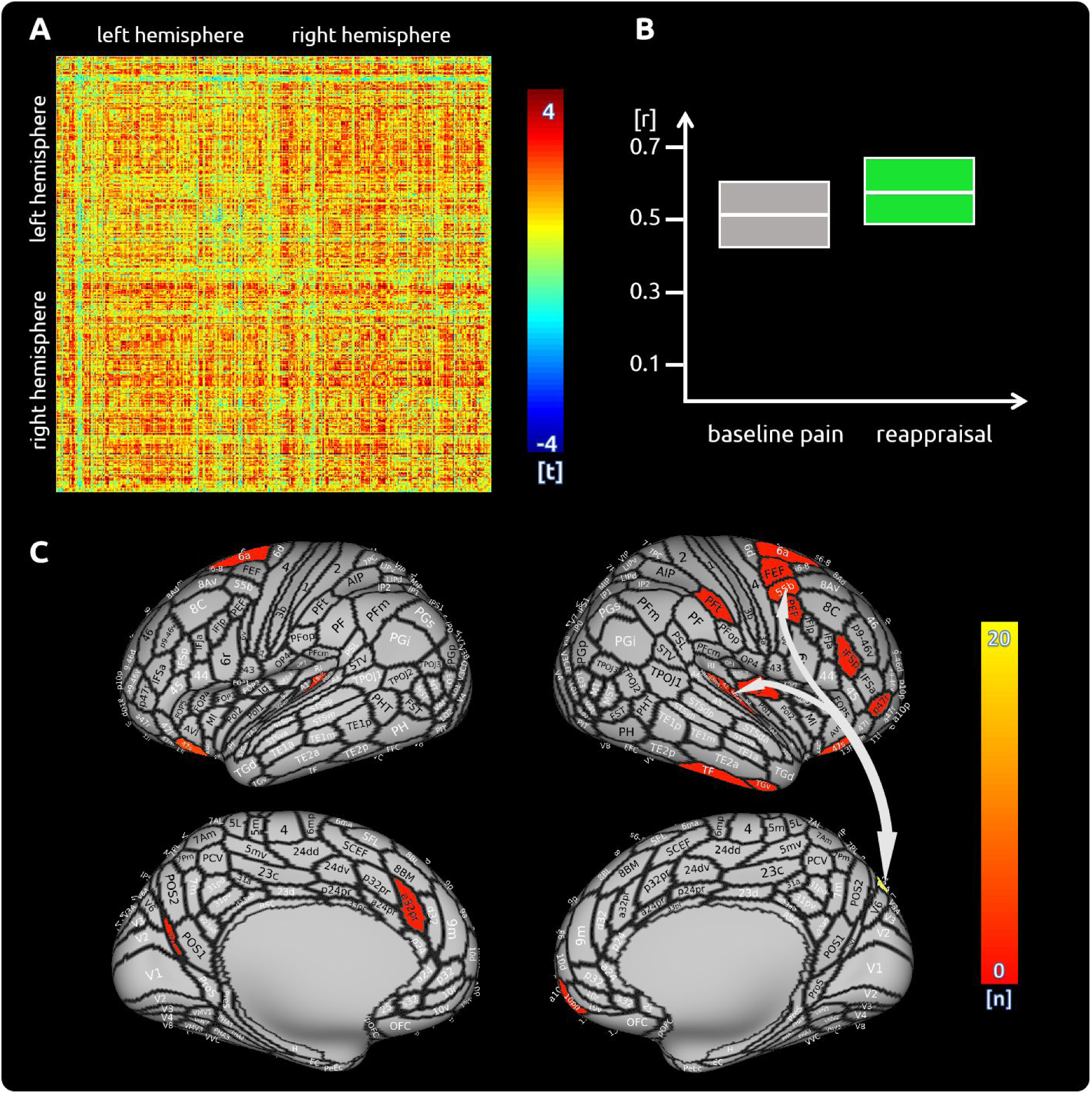
Reappraisal: (A) the confusion matrix shows the statistical results. Each line represents one of the 371 ROIs. The values are mirrored along the principal diagonal of the matrix. A single red dot represents the varying connectivity between two specific brain regions and indicates that a stronger cortical connectivity in a single trial is related to a decrease in pain perception (performance encoding). These findings are the result of the higher connectivity in the trials of the reappraisal task compared to unmodulated pain trials. (B) data from the confusion matrix averaged across all subjects, connections and trials (mean ± standard deviation; for illustration purposes only). (C) Depiction of the cortical regions as defined by the Glasser parcellation. The arrows show the best connected regions; the region V6A in the parieto-occipital cortex has the most connections where connectivity changes are shown to significantly modulate pain intensity. Only regions with at least 3 significant connections (n>2) are included in the cortical map. For more detailed information on the exact connections see the supplementary spreadsheets.

### (4) Conjunction analysis

We did not find any pain-related connectivity changes present in all three conditions.

## DISCUSSION

Here, we aimed to explore how functional and structural connections in the brain contribute to executing cognitive tasks that attenuate pain (Devine and Spanos, 1990; Schulz et al., 2019) by utilising a single-trial analysis approach afforded by ultra-high field imaging. Across three experimental conditions, 20 healthy participants were asked to (a) count backwards, (b) imagine a safe and happy place, and (c) apply a cognitive reappraisal strategy. All strategies resulted in significant pain relief when compared to the unmodulated pain condition. We applied a whole-brain approach on the basis of brain parcellation definitions (Glasser et al., 2016) and explored connectivity patterns during single attempts to attenuate pain. We further explored whether functional connections are facilitated by axonal fibre connections, measured with DTI. All functional connections are discussed in light of our recent findings on BOLD activity (Schulz et al., 2019).

Across all cognitive interventions, our results revealed an *increase* of connectivity pattern throughout the cerebral cortex for all three interventions; a higher functional connectivity was related to particularly effective single attempts to attenuate pain. Therefore, the unmodulated pain trials - which were experienced as considerably more painful - exhibited a lower functional connectivity compared to pain trials during cognitive tasks. This finding has two implications:

*First*, increased connectivity in *task-related* regions is necessary to effectively execute the respective cognitive tasks.

*Second*, contrary to our hypothesis and previous findings, increased connectivity with *pain-related* brain regions (e.g. insular cortex, ACC, or somatosensory cortices) is related to more effective attenuation trials with decreased intensities of pain. These increased *connectivities* are required to actively suppress the *activity* in regions known to contribute to pain processing (Tracey, I., 2008) and are further modulated in the respective task (Schulz et al., 2019). The neuronal activity of these pain-related brain regions are most likely to be actively inhibited, such as by GABAergic neurons in subregions of the insular cortex (Thiaucourt et al., 2018; Watson, 2016), and thus contribute to a lower pain experience by impeding the processing of pain in these insular regions.

### Counting

For the cognitively-demanding counting task, we found a number of well-connected regions that contribute directly or indirectly to the reduction of pain intensity. These regions are located in the parietal and occipito-temporal cortices, overlapping with the modulation of BOLD activity during counting tasks (Johansen-Berg and Matthews, 2002; Schulz et al., 2019). Increased connectivity during counting occurred for connections with pain processing areas, such as divisions of the insular cortex, the posterior cingulate cortex, and the primary and the secondary somatosensory cortices. The highest number of connections to other brain regions during the counting task was found for the right middle insular cortex. Although our analyses do not allow for any assumptions on directionality, the many pain-related functional connectivities between left parietal areas (high BOLD activity) and right insular sub-regions (low BOLD activity) suggest a suppression effect on these insular areas (see Schulz et al., 2019).

Disrupted connectivities during the counting task were observed for the right temporo-parieto-occipital junction (TPJ) to the right posterior insula, as well as to temporo-occipital areas. Given the involvement of the TPJ in attentional processing (Kucyi et al., 2012; Mars et al., 2012), elevated focus on the task may have decreased the transmission during task execution but increased the transmission for the unmodulated pain trials (Schulz et al., 2019). The counting trials are further suggested to require a visual support by imagining the numbers in space (Amalric and Dehaene, 2016). Visual areas in the *left* occipito-temporal cortex connect to and suppress right parietal opercular areas. We also found visual support located in the *right* occipito-temporal cortex that is functionally connected to parietal areas, which in turn suppress the activity in insular sub-regions.

### Safe place

Similar to the counting condition, we found regions in the left and right parieto-occipital cortex to be highly connected to other brain regions. Notably, the parietal cortex is functionally connected without a rise of regional BOLD activity (see Schulz et al., 2019). This effect shows that brain regions can play an important role in pain processing via an exchange of information, where low-scale modulations of cortical activity are not causing large metabolic effects. Moreover, the strong connectivity pattern between left parietal and right insular sub-regions suggests an active suppression of these insular regions initiated by the parietal cortex (as reflected by *increased* functional connectivity between these regions).

We found well-connected regions in the precentral gyrus: area 55b has been shown to be active during listening to stories in the language task of the Human Connectome Project dataset (Glasser et al., 2016). Therefore, the increased connectivity in area 55b may be related to the narrative aspects of the imaginary task in which the participants may recall being actively involved in an event of pleasure and happiness. The premotor and motor areas in the precentral gyrus in particular may reflect the motor aspect of the imagination task (Szameitat et al., 2007; Xie et al., 2015). They are connected to orbitofrontal areas which are thought to initiate top-down pain suppression of ascending pathways (Bingel et al., 2007; Eippert et al., 2009).

For the safe place condition only, we found that the ability to functionally utilise certain pathways is mediated by the strength of axonal fibre connections. These anatomical characteristics are suggested to help the participants with stronger fibre connections to better attenuate pain. This applies especially to connections between middle frontal regions (IFSp and 8C) to the secondary somatosensory cortex (SII). Further functional connections that are supported by the strength of fibre connections project from the frontal cortex to memory-related limbic areas (presubiculum of the hippocampus, entorhinal cortex), which could facilitate memory retrieval for the imagination of pleasant and complex scenes (Braskie et al., 2009; Dalton and Maguire, 2017; Hodgetts et al., 2017; Montchal et al., 2019; Schultz et al., 2015).

### Reappraisal

The best connected region during cognitive reappraisal is located in the higher order visual cortex, area V6A, which is mainly interconnected with several insular and frontal premotor areas. Area V6A is known to contribute to spatial object localisation; a study on monkeys shows that V6A cells are active when executing reaching movements independent of visual or oculomotor processing (Fattori et al., 2005). These cells have also been found to encode body-centred spatial localisation (Hadjidimitrakis et al., 2014). The use of V6A and its connection to other brain areas could help the participants - as required by the task - to focus on the stimulated body site. However, this focussing should be considered as a prerequisite and does not necessarily imply any pain attenuation. Yet the focus on pain has been shown to increase pain perception and pain-related cortical activity (Bantick et al., 2002; Hauck et al., 2007; Peyron et al., 1999). Therefore, as found in the present investigation, the connections from the inferior frontal cortex, the anterior cingulate cortex, the frontal pole, and orbitofrontal cortex are additionally required to utilise cognitive reappraisal (Buhle et al., 2013; Tracey, Irene, 2010) in order to ultimately attenuate the experience of pain (Schulz et al., 2019).

### Analysing pain-related functional connections in the human brain

Unlike previous studies, we almost exclusively found a lower functional connectivity for trials and conditions of higher pain intensity, which could be caused by differences in experimental design and analysis strategies.

In neuroimaging, functional connectivity is considered a joint phase-locked oscillation of spatially distinct cortical regions. Task-based connectivity analyses predominantly utilise a seed-based approach to determine the functional connectivity between a predefined seed region and one or more distant brain regions; such analyses can only take into account the short period during which a task is being executed. However, exact connectivity measures between brain regions would require a sufficient number of samples to quantify the joint in-phase increases and decreases of the BOLD response. In order to estimate a reliable measure of connectivity, we applied a relatively long time window (∼30s) for inflicting pain, for executing the cognitive task, and for reliably determining the connectivity of a single trial. The extended stimulation and the restriction of the analysis to the plateau phase resulted in a solid data basis (330 time points for each subject for each condition) and makes the present investigation more reliable than any other previous work. For instance, a study by Villemure & Bushnell (Villemure and Bushnell, 2009) sampled every 4s but analysed a relatively short time window of 5s painful stimulation to investigate connectivity. Another study analysed just a single data point per trial (3s analysis window, sampling of 3s) before nociceptive laser stimulation to predict pain intensity (Ploner et al., 2010). A further study used 3 data points per trial for connectivity analyses of an experiment in which the pain stimulations lasted 10s (Sprenger et al., 2015). A repeated stimulation at the frequency of the recording (application of 5 brief laser pain stimuli every 3s sampled with a TR of 3s) makes it difficult to separate the connectivity aspects from the general increase of the BOLD response (Ploner et al., 2011).

Furthermore, we used an elaborate artefact cleaning which is recommended for connectivity analyses, particularly for the analysis of pain where a strong and brief painful stimulation can induce task-related movement due to little muscle twitches. Our focus on the analysis of the stimulation plateau makes the connectivity estimates unaffected by the statistical design matrix.

Therefore, the different methodological approaches might have caused our findings to contradict previous studies, in which high levels of pain were shown to increase cortical connectivity between pain processing brain regions (Sprenger et al., 2015). Villemure & Bushnell (Villemure and Bushnell, 2009) and Ploner et al. (Ploner et al., 2011) found a stronger connectivity in pain processing brain regions for conditions that increased the intensity of pain (i.e. increased attention, more negative emotion). The connectivity of the inferior frontal cortex for an emotional condition, and the connectivity of the superior parietal cortex, and the entorhinal cortex for the attentional condition were found to modulate cortical processes (Villemure and Bushnell, 2009).

Other studies investigated the connectivity in the descending pain control system and observed an increase of connectivity between the perigenual ACC and the PAG during a pain-relieving placebo intervention (Eippert et al., 2009). Given the lower signal-to-noise ratio in mid-brain areas, this finding could not be replicated in any of the present conditions with the current whole-brain approach and a strict correction for multiple comparisons (Winkler et al., 2014). By lowering the statistical threshold, we found a modulation of pain intensity-dependent functional connectivity from the PAG to regions that contribute to pain processing, such as the anterior ventral insula (t>2), the midcingulate cortex (t>2.5), and the nucleus accumbens (t>3), indicating a stronger connectivity for the pain condition with cognitive modulation.

Further studies directly investigated the functional connectivity in the brain in response to different intensities of pain stimuli. Sprenger et al. (2015) found an increase of connectivity in subcortical nuclei for the higher of two pain conditions. Similarly, an increased connectivity has been found in response to cold pain stimulation. The authors reported a significant correlation across the entire time course of the experiment between predefined regions that are known to be involved in the processing of pain(Wilcox et al., 2015). As discussed above, our data showed that the decrease of pain is predominantly related to an increase of cortical connectivity in both *pain-related* regions (e.g. subregions of the insular cortex) and *task-related* brain regions (subregions in the frontal and parietal cortex).

### Summary

The present investigation resembles a clinical intervention in which a pain patient would be taught to utilise cognitive strategies to attenuate pain. Here, we investigated which cortical connection contributes to particularly effective trials to attenuate pain. In contrast to previous research, we revealed an increased connectivity for the single attempts that resulted in decreased pain. This applies to the classical pain processing regions (e.g. insula, cingulate cortex, and somatosensory cortices). Although we found different connectivity patterns for all interventions, the general mechanism was universally valid. The present findings are suggested to open a new window in the understanding of cortical processes that are associated with high levels of long-lasting tonic or chronic pain. As a consequence, clinical treatments that would aim to decrease cortico-cortical connections are suggested to have a rather detrimental effect on pain relief in patients suffering from chronic pain. Therefore, future studies would be needed to investigate the effect of cognitive interventions on the intracortical connectivity in pain patients.

## METHODS

Twenty two healthy human subjects (18 female/4 male) with a mean age of 27±5 years (21 - 37 years) participated in the experiment. Two of the female subjects were excluded as a result of insufficient data quality. All subjects gave written informed consent. The study was approved by the Medical Sciences Interdivisional Research Ethics Committee of the University of Oxford and conducted in conformity with the Declaration of Helsinki.

The experiment, the participants, and the behavioural data have been described in detail in our previous publication (Schulz et al., 2019). The task consisted of four conditions (see table 1) across 4 separate blocks, where each block comprised of 12 trials from the same condition. The first condition was always the unmodulated condition; the order of the conditions with cognitive interventions were randomised using Matlab (randperm). In all conditions and trials the subjects received cold pain stimuli on the dorsum of their left hand delivered by a thermode (Pathway II; Medoc Ltd, Ramat Yishai, Israel). The subjects were prompted to rate pain intensity and pain unpleasantness. A numerical and a visual analogue scale (VAS), ranged between 0 - 100 in steps of 5 points, was used to assess the pain ratings. The endpoints of the scale were determined as no pain (0) and the maximum pain the subjects were willing to tolerate (100). Single trial ratings were recorded after each trial.

**Table 1:** Conditions and Instructions.

|  |  |
| --- | --- |
| (0) pain, non-modulated | Concentrate only on the pain. |
| (A) attentional shift | Count backwards from 1000 by sevens. |
| (B) imaginal strategy | Imagine that you are in a safe and happy place that you know very well. That place has the colours you like and you hear the music you like. There are only people around that you want to have around you. You feel well and comfortable. |
| (C) cognitive reappraisal | Concentrate on the cool and tingling sensations in your arm and reinterpret these sensations as not painful. |

The thermode temperature for painful stimulation for each subject was determined in an extensive practise session one week prior to scanning and was individually adapted to a VAS score of 50. The 40s of painful stimulation were then preceded by a rest period of 10s at 38°C thermode temperature. The first 10s were not included in the analysis. The mean temperature of cold pain application across subjects was 7°C with a standard deviation of 3.6°C. In order to avoid habituation effects, the thermode temperature during painful stimulation was oscillating with ⅛ Hz at ± 3°C (Lautenbacher et al., 1995; Stankewitz et al., 2013).

### Data Acquisition

Imaging data were acquired on a 7T Siemens MRI scanner. Each volume comprised 34 axial slices of 2 mm thickness and 2 × 2 mm in-plane resolution with 1mm gap between slices. The repetition time (TR) was 1.96s, the echo time (TE) was 25ms (flip angle 90°), the field of view (FOV) was 220 × 220 mm, and the matrix size was 110 × 110 pixels. A T1-weighted structural image (isotropic 1mm^3^ voxel) was acquired for the registration of the functional images to the MNI (Montreal Neurological Institute) template. Two sequences of diffusion tensor images (DTI) were recorded with L>>R and R>>L phase encoding direction. 64 directions were recorded with a TR of 9.3s, a TE of 63ms, and an acceleration factor of 2. The length of the edge of the isotropic voxels was 1.2 mm.

### Image processing - preprocessing of functional connectivity data

The data were preprocessed with FSL (Jenkinson et al., 2012). The preprocessing of the *functional data* consisted of brain extraction, high-pass filtering with a frequency cutoff of 1/90 Hz, a spatial normalisation to the MNI template, a correction for head motion during scanning registered to the MNI template, and a spatial smoothing (6 mm FWHM). The data were further semi-automatically cleaned of artefacts with independent component analysis (ICA) (Griffanti et al., 2014; Salimi-Khorshidi et al., 2014). The number of components had been set *a priori* to 200. Artefact-related components were removed from the data. The design matrix for painful stimulation, including the temporal derivative, were then regressed out from the data in Matlab (The Mathworks, USA).

### Image processing - preprocessing of structural connectivity data

Preprocessing of DTI data was performed using FSL. FSL preprocessing included (i) correcting susceptibility induced distortions (“topup”), (ii) skull stripping (“bet”), and (iii) corrections for eddy currents and head motion (“eddy”). We finally (iiii) determined the strength of structural connectivity between cortical regions (“bedpostx” and “probtrackx”) defined by the Glasser atlas. For tractography, we used the “one-way condition”, with 5000 samples, 2000 steps per sample, a step length of 0.5, and a fibre threshold of 0.1. The number of streamlines quantifies the strength of structural connectivity for two brain regions of a subject.

### Image processing - extraction of regions of interest data

The time series of functional volumes were converted to MNI space and subsequently projected to surface space by using the “Connectome Workbench” package. We used a template that allowed to project from 3D standard MNI space to 2D surface space. Regions of interest (ROIs) were defined by subdividing the cortical surface into 180 regions per hemisphere (Glasser et al., 2016). Six further regions (5 bilateral) that are important for the processing of pain, such as the PAG, the thalamus and the amygdala, were also included. Latter ROIs were based on the Oxford Atlas, implemented in FSL.

### Image processing - computation of single trial functional connectivity scores

The time courses for all voxels of cortical activity for a specific region of the Glasser Atlas, e.g. the middle insula, were extracted. We computed principal component analyses (PCA) separately for each ROI and subject and selected the first component (Matlab, The MathWorks, Inc., USA). The plateau phase of the last ∼30s of painful stimulation (15 data points) has been extracted from each region and trial for each subject and condition. Outliers were removed from the data by using the Grubbs’ test (Grubbs, 1950). These 15 data points determined the connectivity for a brain region for a given trial. Correlation coefficients were computed for each trial and for each of the 371 ROIs with the remaining 370 ROIs. The single trial correlation coefficients were Fisher Z-transformed and fed into group-level statistical analysis.

### Image processing - structural connectivity data

DTI data were also analysed in FSL. The processing steps included a median filter, a correction for susceptibility distortions, and fibre tracking from the same aforementioned brain regions (Glasser parcellation - see above).

### Statistical modelling

The statistical analysis for the connectivity between cortical regions has been performed in Matlab. To explore the relationship of fluctuating cortical connectivity and the variable pain experience, we computed linear mixed effects models (LMEs) that related the single trial correlation coefficients between two brain regions to the pain intensity scores (Michail et al., 2016; Schulz et al., 2015). Each condition in the model included the data for the respective intervention plus the trials of the unmodulated pain condition. Using the Wilkinson notation (Wilkinson and Rogers, 1973), the model can be expressed as:

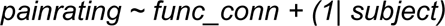

The included fixed effects (*func_conn*) essentially describe the magnitudes of the population common intercept and the population common slope for the relationship between cortical connectivity and pain perception. The included random effects (1| subject) are used to model the specific intercept differences for each subject.

This single trial approach has two major advantages: *First*, it takes the within-subjects variable performance of the pain attenuation attempts into account; e.g. a more successful attempt to attenuate pain (as reflected by lower pain ratings) is considered to cause a stronger cortical effect (compared to the unmodulated pain condition) than a less successful attempt. *Second*, it also takes into account the more natural fluctuation of the unmodulated pain trials (without cognitive intervention).

We further analysed whether individual differences in functional connectivity could be explained by individual structural characteristics of the brain. In other words, we analysed whether the functional connectivity that leads to a single subject’s successful pain attenuation is facilitated by that subject’s high number of fibre tracts. In a similar vein, a poor functional connectivity that is not able to contribute to pain attenuation might be caused by a low number of fibre tracts.

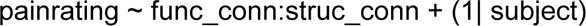

We considered only functional connections with a t-value >2 as potentially modulated by structural connections.

To correct for multiple comparisons we applied a randomisation approach. Behavioural data were shuffled and the entire analysis was repeated 5000 times. The highest absolute t-values of each repetition across the whole confusion matrix was extracted. This procedure resulted in right-skewed distribution for each condition. Based on these distributions, the statistical thresholds were determined using the “palm_datapval” function implemented in PALM (Winkler et al., 2014).

### Funding

● Deutsche Forschungsgemeinschaft
  ○ Enrico Schulz
● Wellcome Trust: 090955/Z/09/Z and 083259/Z/07/Z
  ○ Irene Tracey
● NIHR Oxford Biomedical Research centre
  ○ Irene Tracey
● Medical Research Council: G0700399
  ○ Irene Tracey

The funders had no role in study design, data collection and interpretation, or the decision to submit the work for publication.

### Ethics

Human subjects: Informed consent and consent to publish was obtained in accordance with ethical standards set out by the Declaration of Helsinki (1964) and with procedures approved by the Medical Sciences Interdivisional Research Ethics Committee of the University of Oxford (REC ref: MSD-IDREC- C1-2014-157).

## SUPPLEMENTARY MATERIAL

**Supplementary Figure 1.**
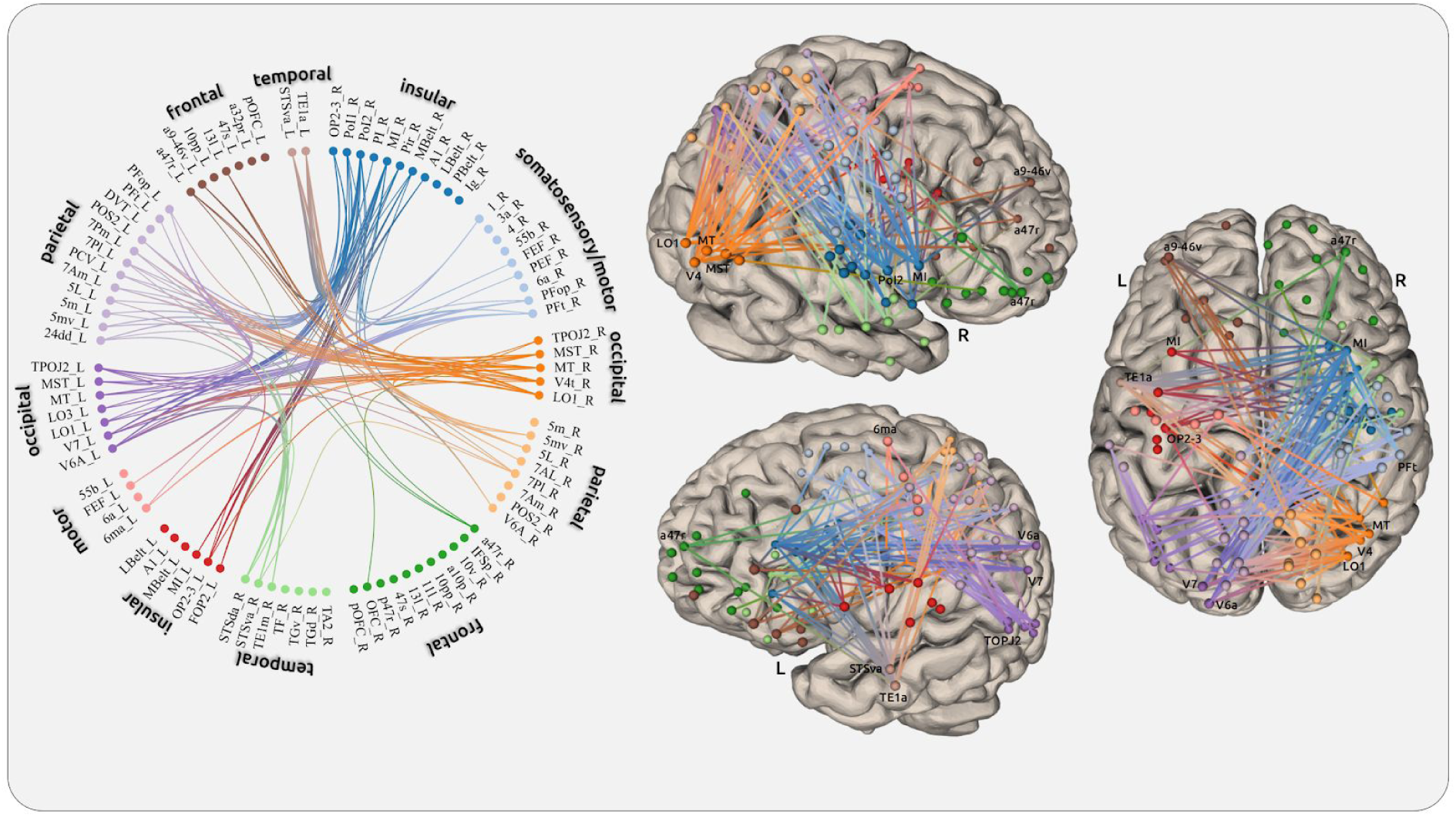
The connectivity plots show the statistical results of the counting condition. We selected 89 regions that showed at least 3 significant connections in any of the 3 conditions. For more detailed information on the exact connections see the supplementary spreadsheets.

**Supplementary Figure 2.**
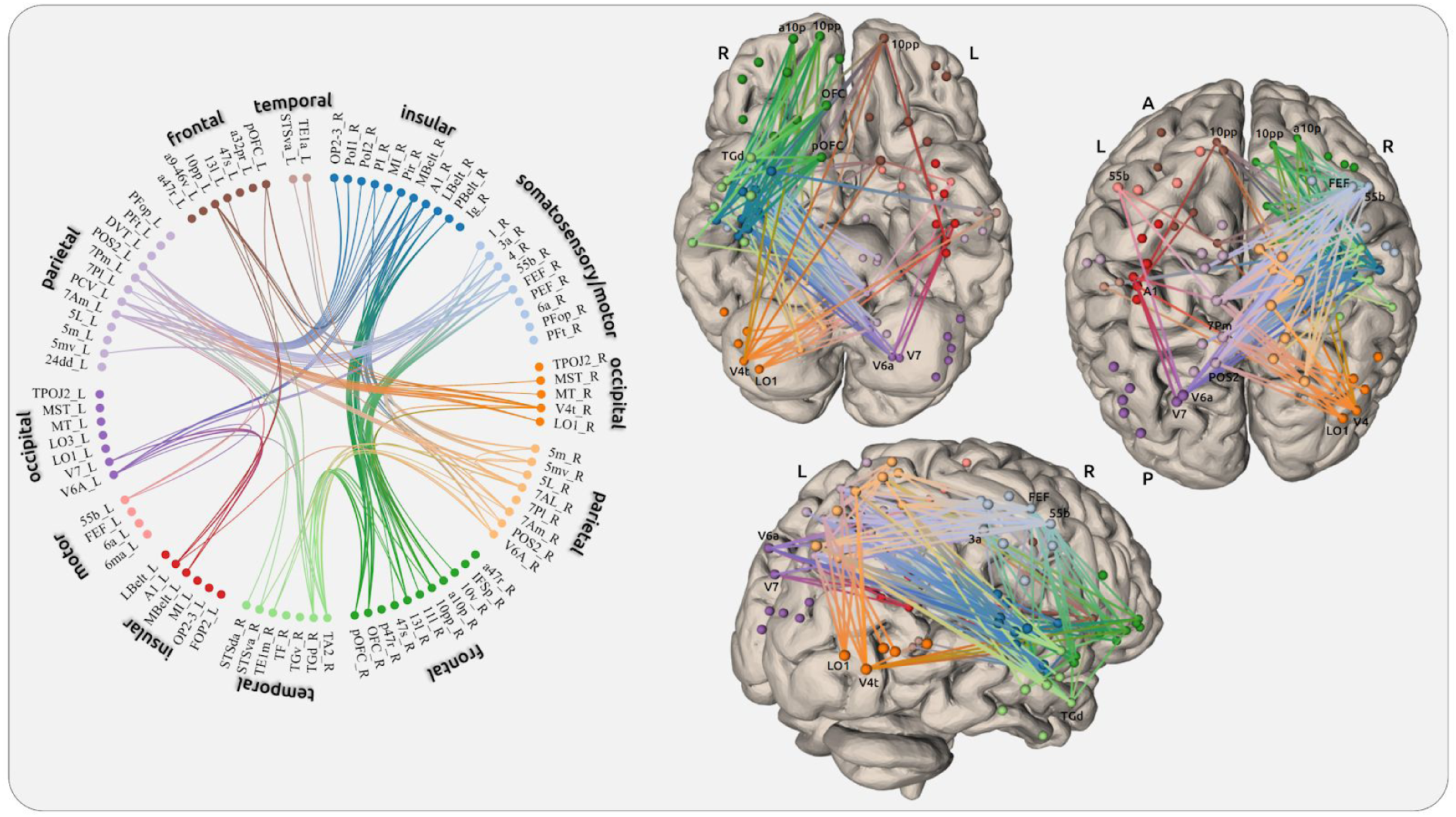
The connectivity plots show the statistical results of the “safe place” condition. We selected 89 regions that showed at least 3 significant connections in any of the 3 conditions. For more detailed information on the exact connections see the supplementary spreadsheets.

**Supplementary Figure 3.**
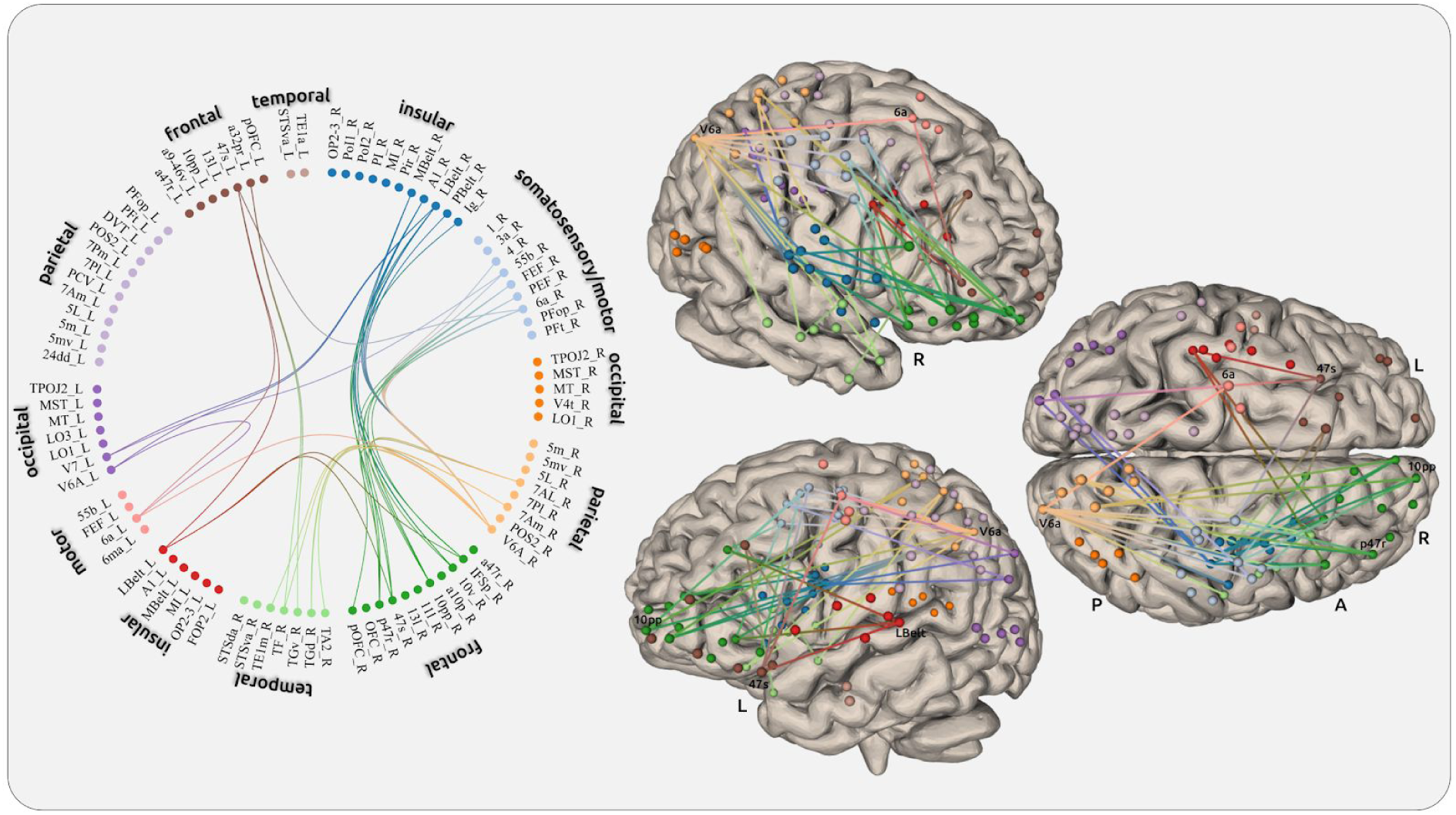
The connectivity plots show the statistical results of the reappraisal condition. We selected 89 regions that showed at least 3 significant connections in any of the 3 conditions. For more detailed information on the exact connections see the supplementary spreadsheets.

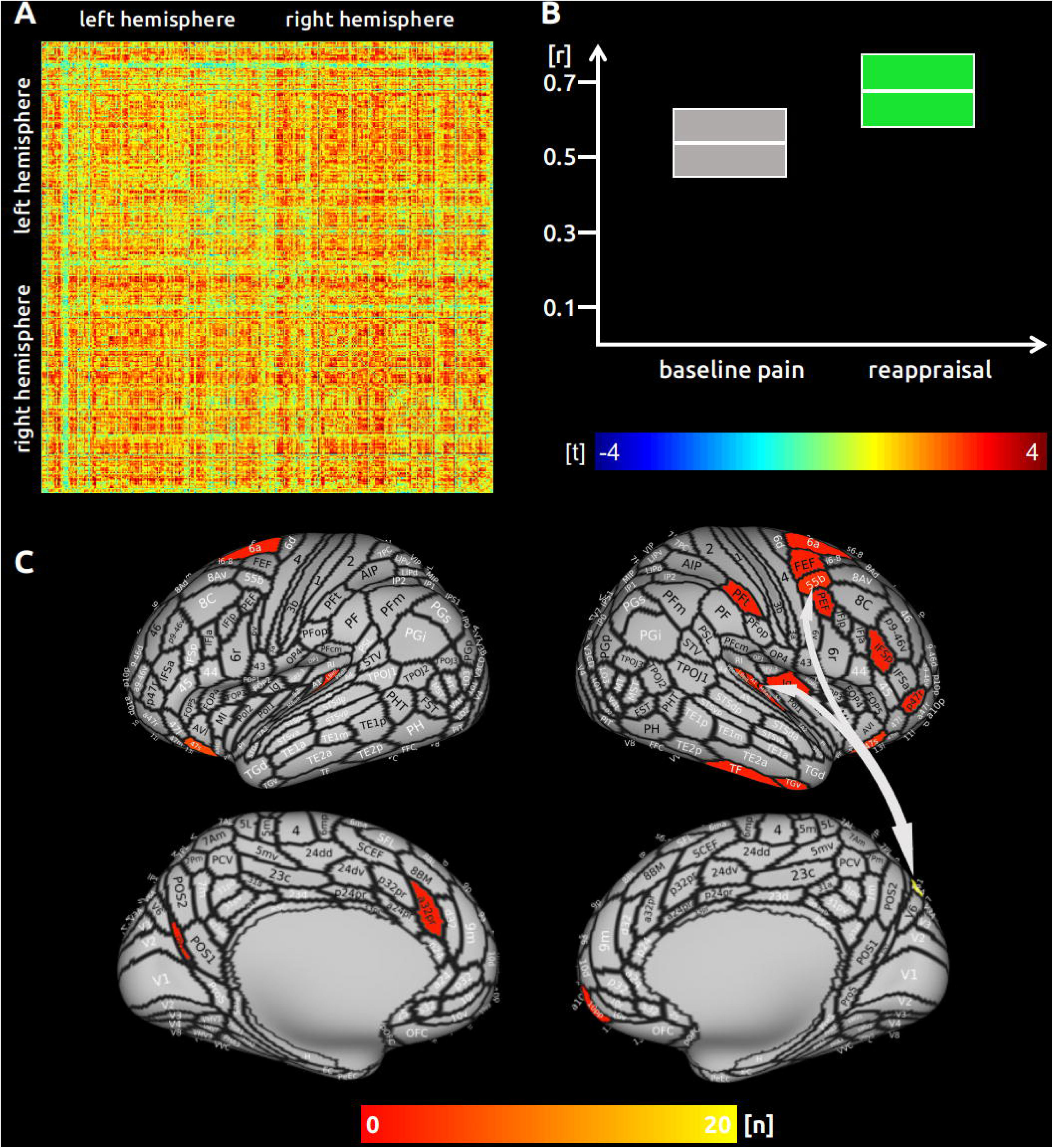

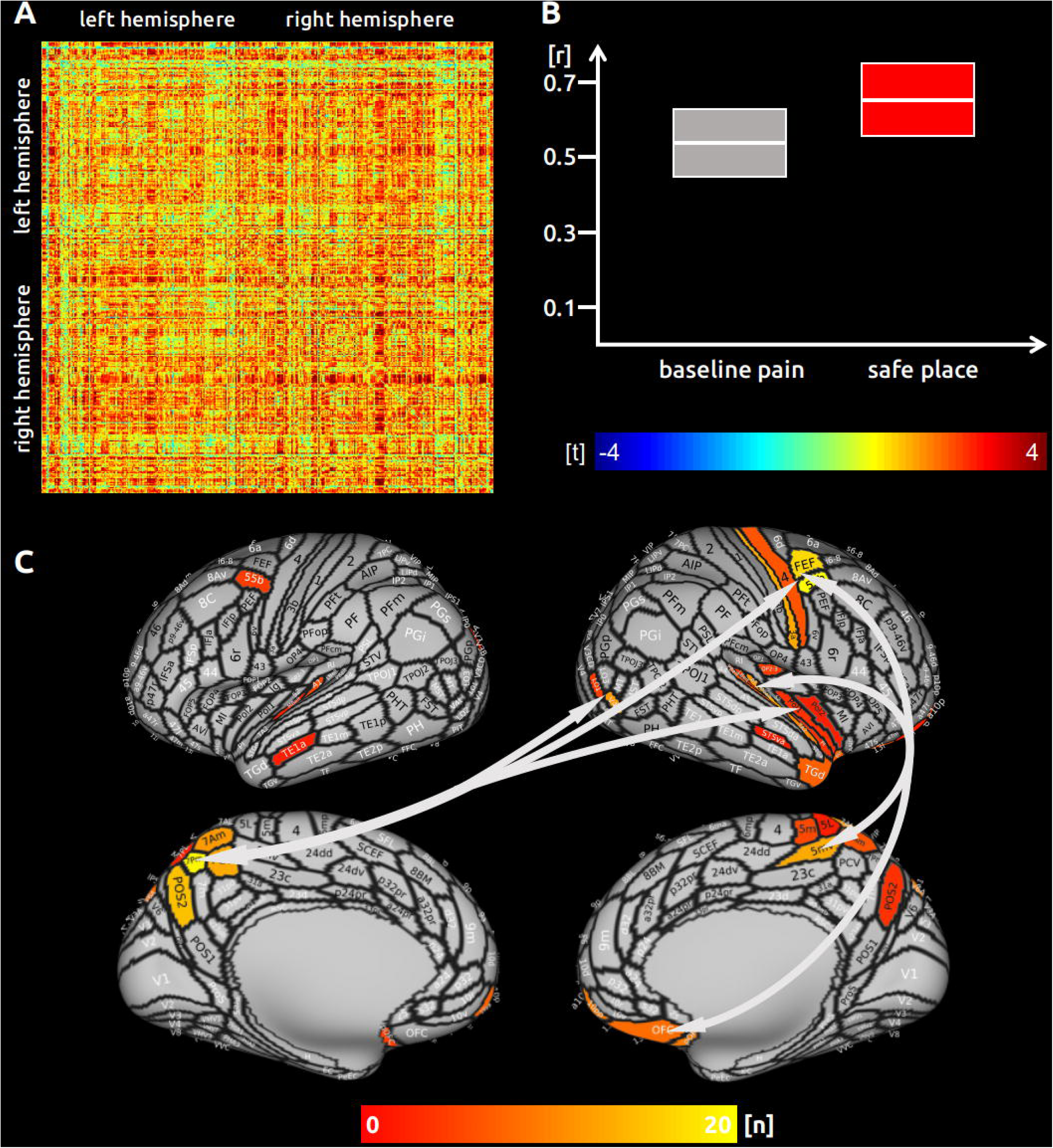

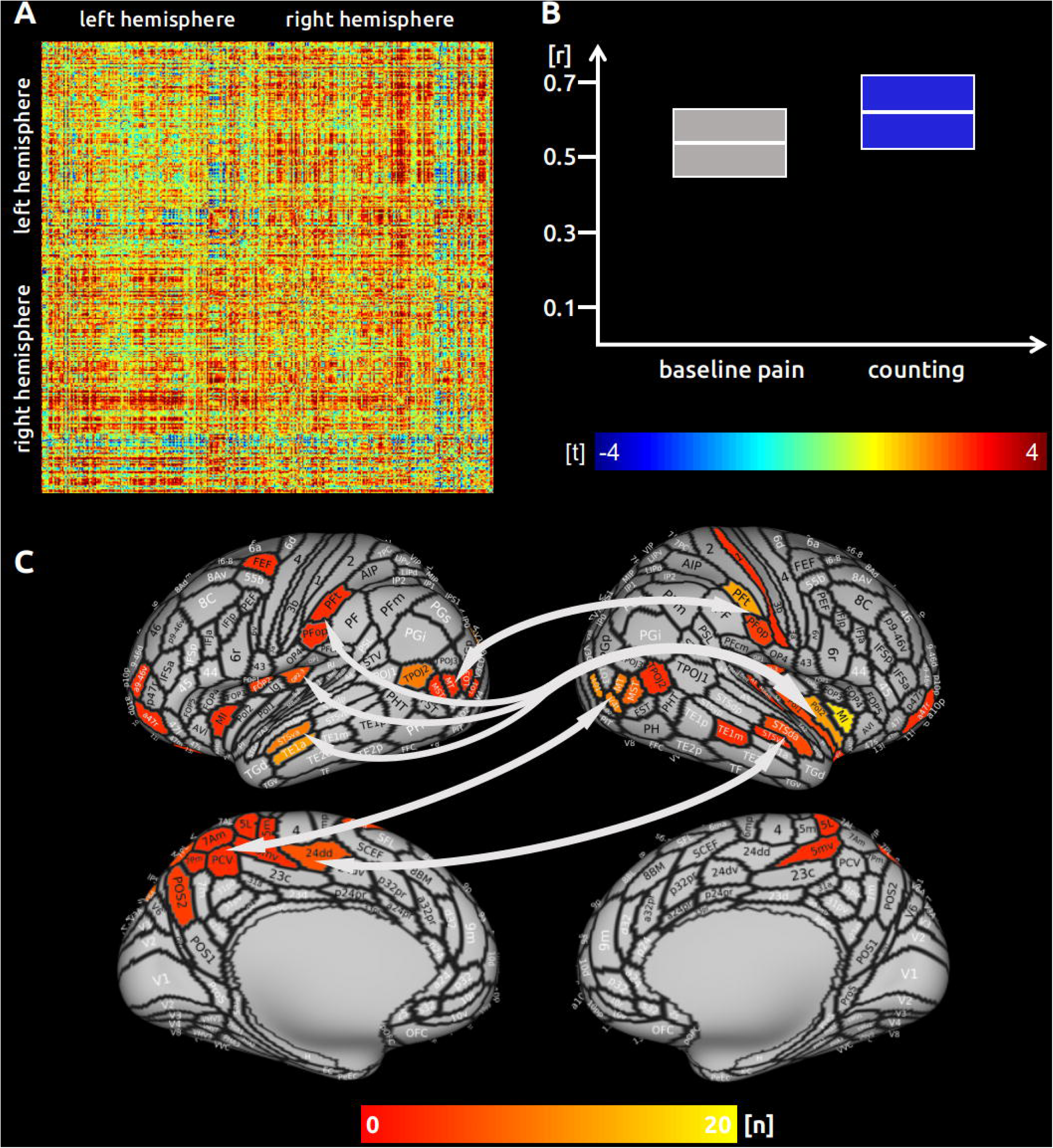

