## Supplementary figures and images for "Ultra-high field imaging reveals increased whole brain connectivity underpins cognitive strategies that attenuate pain"

### Supplementary Figure 1

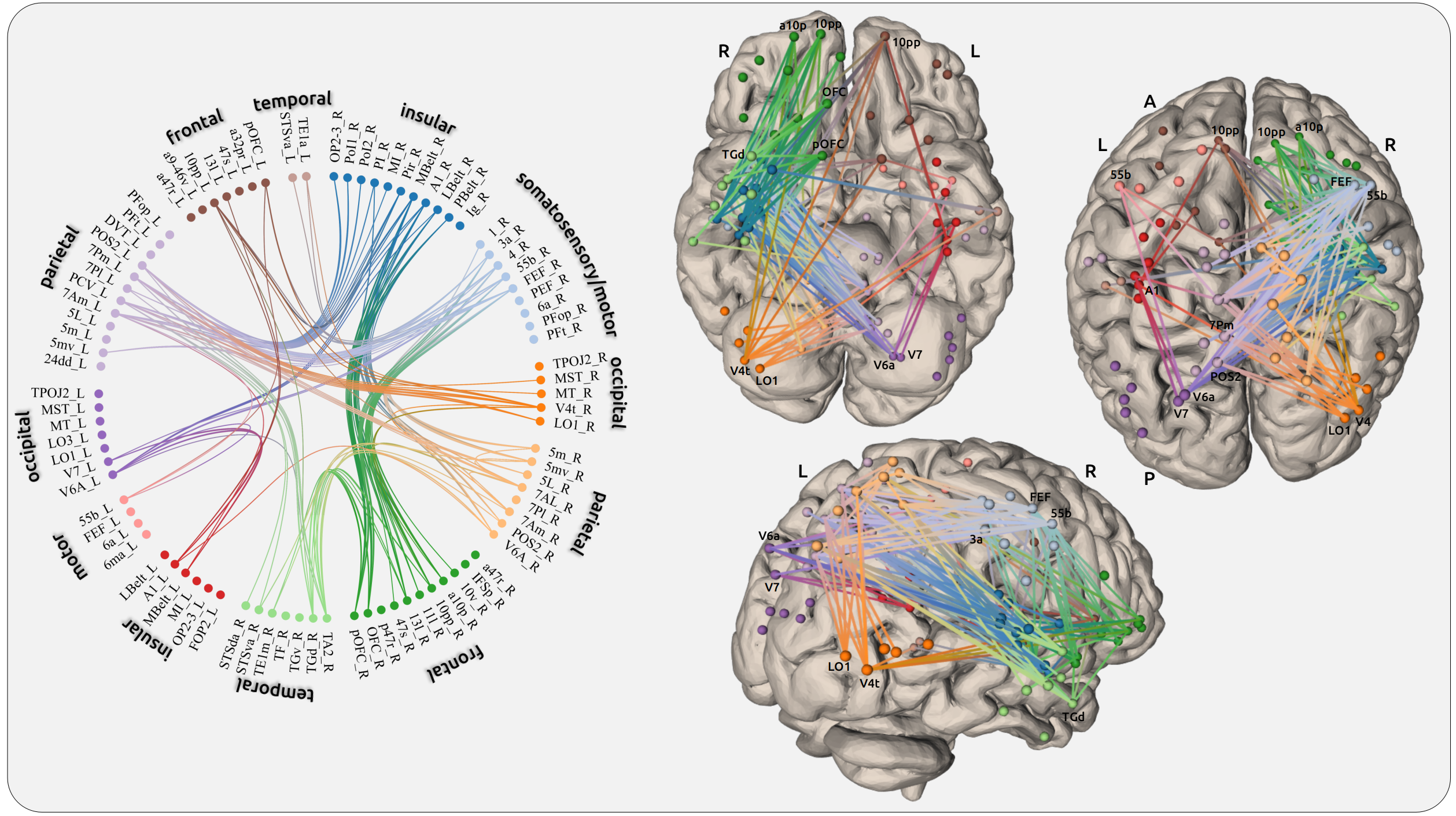

### Supplementary Figure 2

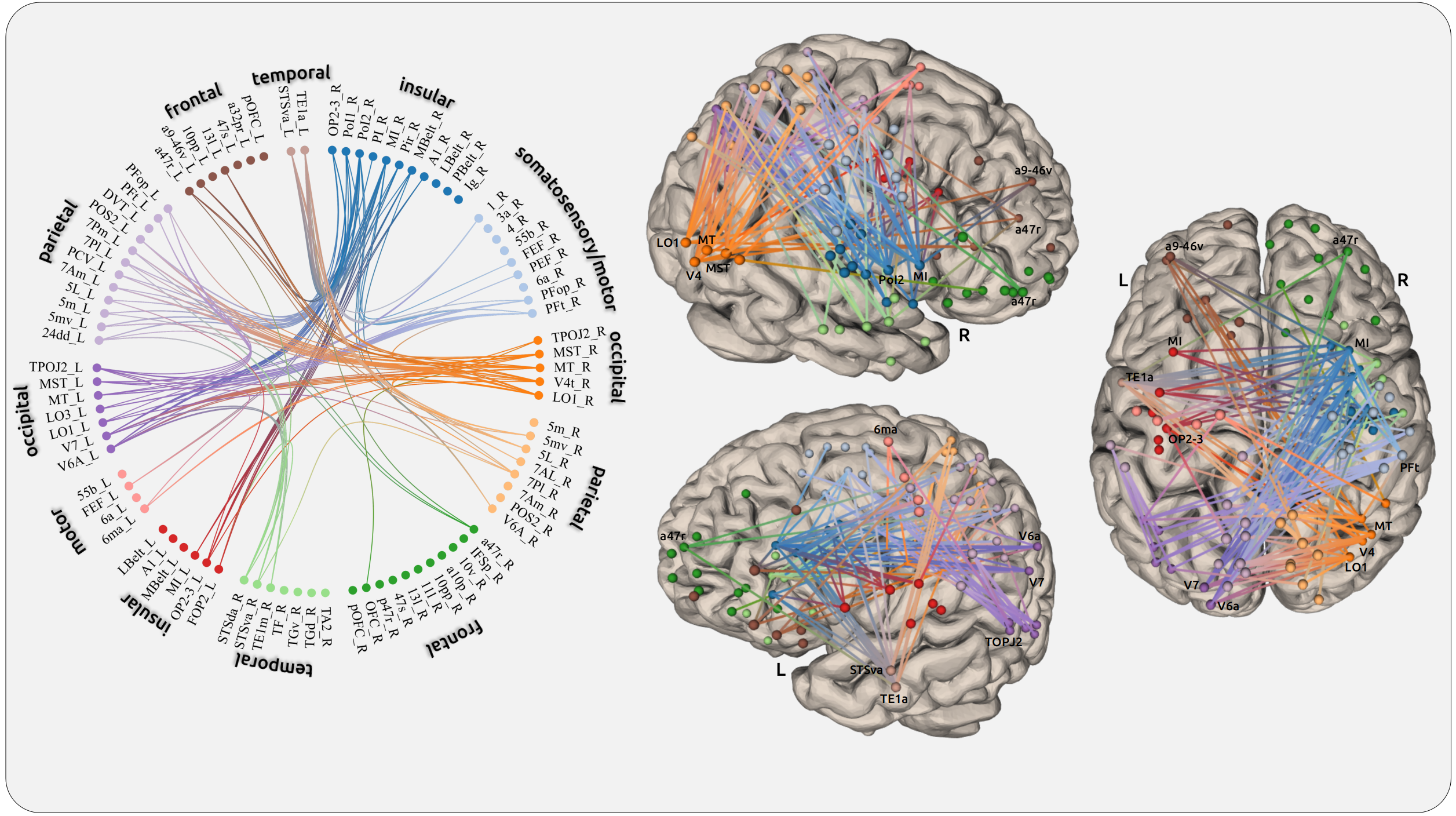

### Supplementary Figure 3

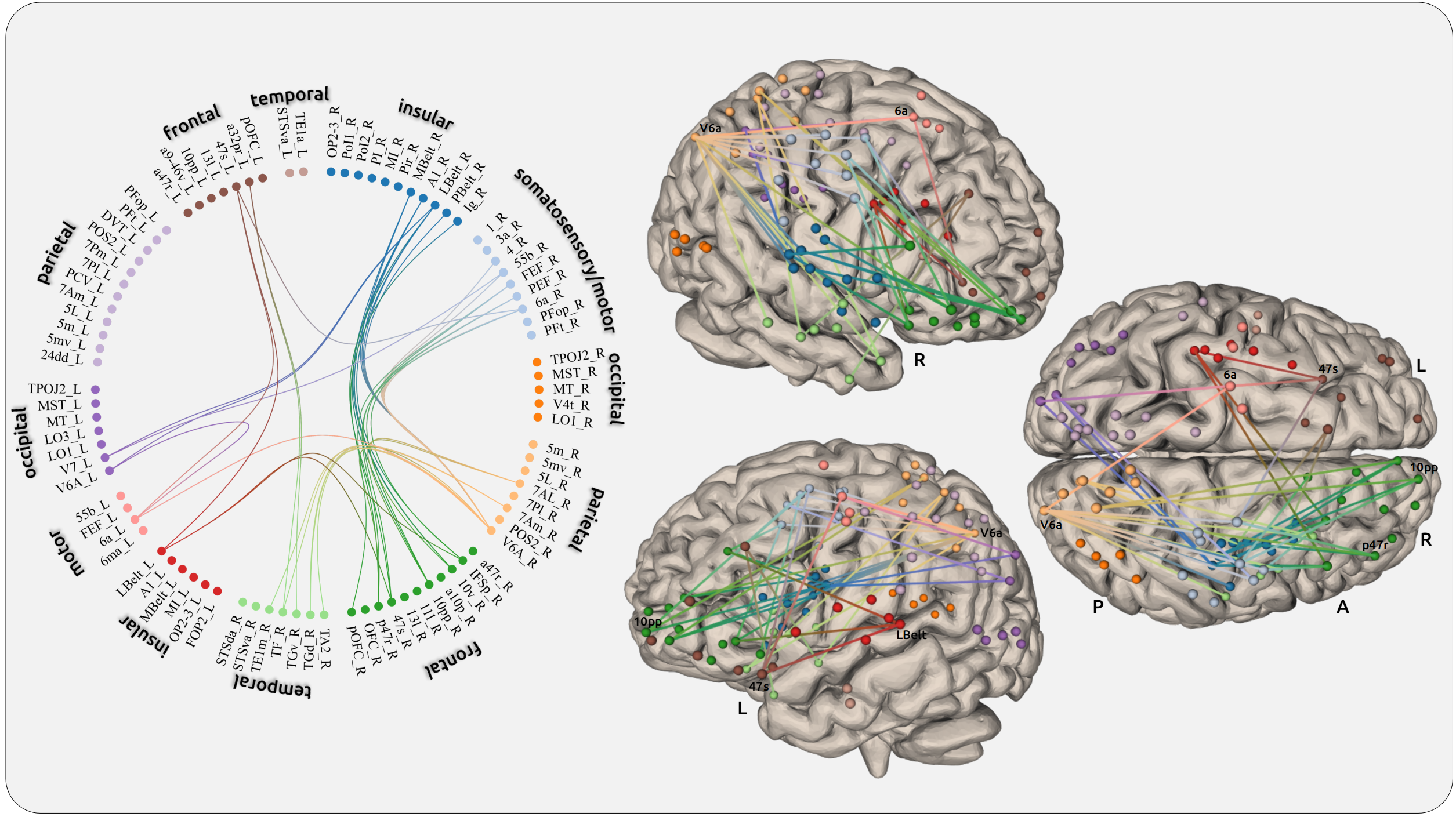
